# Agent-Based Simulation of Cortical Microtubule Band Movement in Arabidopsis Zygotes

**DOI:** 10.1101/2024.10.17.618799

**Authors:** Tomonobu Nonoyama, Zichen Kang, Hikari Matsumoto, Sakumi Nakagawa, Minako Ueda, Satoru Tsugawa

## Abstract

*Arabidopsis thaliana* zygotes form a circumferential cortical microtubule (CMT) band that moves upward along the longitudinal axis of the cylindrical cell; however, how and why this CMT band moves remain unknown. Based on several recently proposed descriptive feedback mechanisms of CMT reorientation, we hypothesized that a directional cue (DC) changes CMT orientation. We constructed an agent-based simulation model with a DC to observe what happens during CMT formation using different parameters. We determined that the output CMT band can become larger than the input DC zone and identified two types of self-organization processes depending on the parameters used. Furthermore, using well-defined parameters that form a static CMT band, we artificially changed the position of the input DC zone, revealing that CMT band movement requires a DC to move at an appropriate speed.

## Introduction

Arabidopsis (*Arabidopsis thaliana*) zygotes are single cells formed by the fusion of egg and sperm cells during plant sexual reproduction. During zygote elongation, the cell adopts a cylindrical shape with a circumferential cortical microtubule (CMT) band near the subapical region; this CMT band maintains a uniform width and moves following the growing tip^1,2^. CMT band formation is critical for establishing the plant apical–basal axis because it is thought to guide the subsequent formation of the first unequal division plane. However, how and why the CMT band moves remain unknown.

As CMTs guide the deposition of cellulose microfibrils and thus, anisotropic cell growth^3,4,5,6^, CMT dynamics are closely linked to cell and tissue morphogenesis. The relationship between the plant cell and CMTs is often associated with mechanical feedback between the stress distribution derived from cell geometry and CMT orientation^7^. CMT-severing proteins cut CMTs perpendicular to stress orientation^8^. Several reports describe the similarity between stress profiles on the cell surface and the distribution of CMT orientation^9,10,11^. Importantly, CMTs display increased polymerization in regions of increased tension^12,13^, suggesting that CMTs may respond to tensile intensity. However, default CMT formation can be overwritten by the addition of artificial tensional cues^14,15^. CMTs return to their original orientation after 10 h, strongly suggesting that CMTs respond not only to the tension intensity itself but also to changes or fluctuations in tension^15,16^. Therefore, CMTs can be modified by changing the intensity or fluctuation of tension within a specific zone.

Based on experimentally obtained CMT–CMT collision probability^17,18^, several agent-based simulations incorporating growth and shrinkage of a single CMT and its interactions have been developed^20,21,22,23,24,25^, including numerical studies implementing growth and/or shrinkage of a single CMT, CMT interactions (zippering, collision-induced catastrophe, and crossover), and nucleation on the existing CMT (branching)^19,20,21^, as well as theoretical studies capturing possible CMT behaviors using mean-field theory^22,23,24,25^. The effect of geometry-induced catastrophe is critical for CMT band formation during embryogenesis^26,27^. In xylem cells, the nucleation rate at specific locations changes, resulting in a CMT pattern with gaps^28,29^. Thus, simulations of CMT patterns mainly modify collision-induced catastrophe, geometry-induced catastrophe, nucleation, and branching rates.

The concept of a directional cue (DC) for CMTs has also been proposed as an external or intrinsic factor that enables CMTs to reorient in response to phenomenological factors, such as mechanical stress, phytohormone signals, or polarity markers (e.g., specific proteins). Briefly, the DC modifies the properties of the CMT, including its nucleation rate, growth rate, and orientational angle within a specific zone, such as the mechanically distinguishable region discussed above. Two possible DCs have been proposed. The first is a DC that results in CMT reorientation^30^, and the second is a dynamic instability (DI) framework at the cellular scale that results in an increased CMT growth rate without CMT reorientation^31^. From the physical perspective of the CMT polymerization process, a compressive force derived from tension perpendicular to the existing CMT promotes an increase in the CMT polymerization rate^12,13^.

In this study, we explored the first concept by performing an agent-based simulation of CMT movement in Arabidopsis zygotes to identify the parameters necessary for the formation of the moving CMT band. We used Arabidopsis zygote data to define a CMT dynamic space (CDS) with the width and movement speed of the moving CMT band. We adopted the reorientation of CMTs as the DC and defined a specific region with a DC as a DC zone. We then reconstructed an agent-based model of CMTs with DCs, with the aim of extracting the model parameter range that best explains the width and movement speed of the CMT band. We analyzed which parameters modify the resulting width of the CMT band. Finally, we systematically modified the movement speed of the DC zone and found that it should be restricted within a certain range in order to be consistent with the observed data; otherwise, movement of the CMT band lags behind the movement of the DC input speed. Our simulation suggests that an appropriate speed of movement of the DC zone is required to form a moving CMT band.

## Methods

To understand the CMT dynamics beneath the membrane of the Arabidopsis zygote, an agent-based simulation was developed focusing on the cylindrical cell geometry during the zygote elongation stage. The agent-based model incorporates the growth, shrinkage, and interaction events between agents (CMTs), including zippering, crossovers, and induced catastrophes. In addition, a DC was implemented, which modifies the angle of the newly added CMT plus end. The effect of the finite tubulin pool that restricts the total length of the CMT was also taken into account.

### Agent-based model of CMTs on the cylindrical cell surface

The agent-based model of the CMTs was based on previous work^27,30^, as shown in Fig. 1. There are six crucial parameters of CMT dynamics: (1) the growth speed of the plus end, *v*^+^ (Fig. 1A); (2) the shrinkage speed of the plus end, *v*^−^ (Fig. 1B); (3) the shrinkage speed of the minus end (treadmilling speed initiated from nucleation), *v*^*tm* 32,33^; (4) the rate of change from shrinkage to growth at the plus end (rescue rate), *r*_*r*_; (5) the rate of change from shrinkage to growth at the plus end (spontaneous catastrophe rate), *r*_*c*_; and (6) the nucleation rate of CMTs, *r*_*n*_. The parameter values are summarized in Table 1. In most simulations, cylindrical cell geometry was used with a radius *R* and height *H*. The height coordinate on the cylinder is denoted by *z*. CMT dynamics were simulated on the cylindrical surface with boundaries at the top and bottom; CMTs beyond the boundary were not taken into account. The simulation was iterated *N* times with the time interval *Δt* for each step. The simulation time interval *Δt* was 0.2 s. Most of the iterations in simulations were 12,000 steps, corresponding to a total of approximately 40 min.

**Table 1.** Simulation parameters and variables with their default values. The height and radius of the cylinders in the simulation are based on observed data from Arabidopsis zygotes^34^. The threshold angle of zippering and the probabilities of induced catastrophes and crossovers were based on previous work^17, 25^. The strength of the directional cue (DC) was based on previous work^30^. CMT, cortical microtubule.

| Parameter | Value | Meaning |
| --- | --- | --- |
| $H$ | 20 ($\mu\text{m}$) | Height of the cylinder |
| $R$ | 5.0–15.0 ($\mu\text{m}$) | Radius of the cylinder |
| $N$ | 6,000; 12,000 | Simulation steps |
| $\Delta t$ | 0.02 (s) | Time interval of the simulation steps |
| $v^+$ | 0.08 ($\mu\text{m/s}$) | Growth speed of the plus end |
| $v^-$ | 0.16 ($\mu\text{m/s}$) | Shrinkage speed of the plus end |
| $v^{tm}$ | 0.01 ($\mu\text{m/s}$) | Shrinkage speed of the minus end |
| $r_r$ | 0.007 (1/s) | Rescue rate |
| $r_c$ | 0.001–0.01 (1/s) | Catastrophe rate |
| $r_n$ | 0.001 (1/s) | Nucleation rate |
| $\theta^*$ | 40 ($^\circ$) | Threshold angle of CMT interactions |
| $P_{cat}$ | 0.5 | Probability of induced catastrophe |
| $P_{cro}$ | 0.5 | Probability of crossover |
| $b_d$ | 0.0001–0.01 | Strength of the DC |
| $\mu_T$ | 10.0 ($\mu\text{m}$) | Position of the center of the DC zone |
| $\sigma_T$ | 2–10 ($\mu\text{m}$) | Width of the DC zone |

**Fig. 1.**
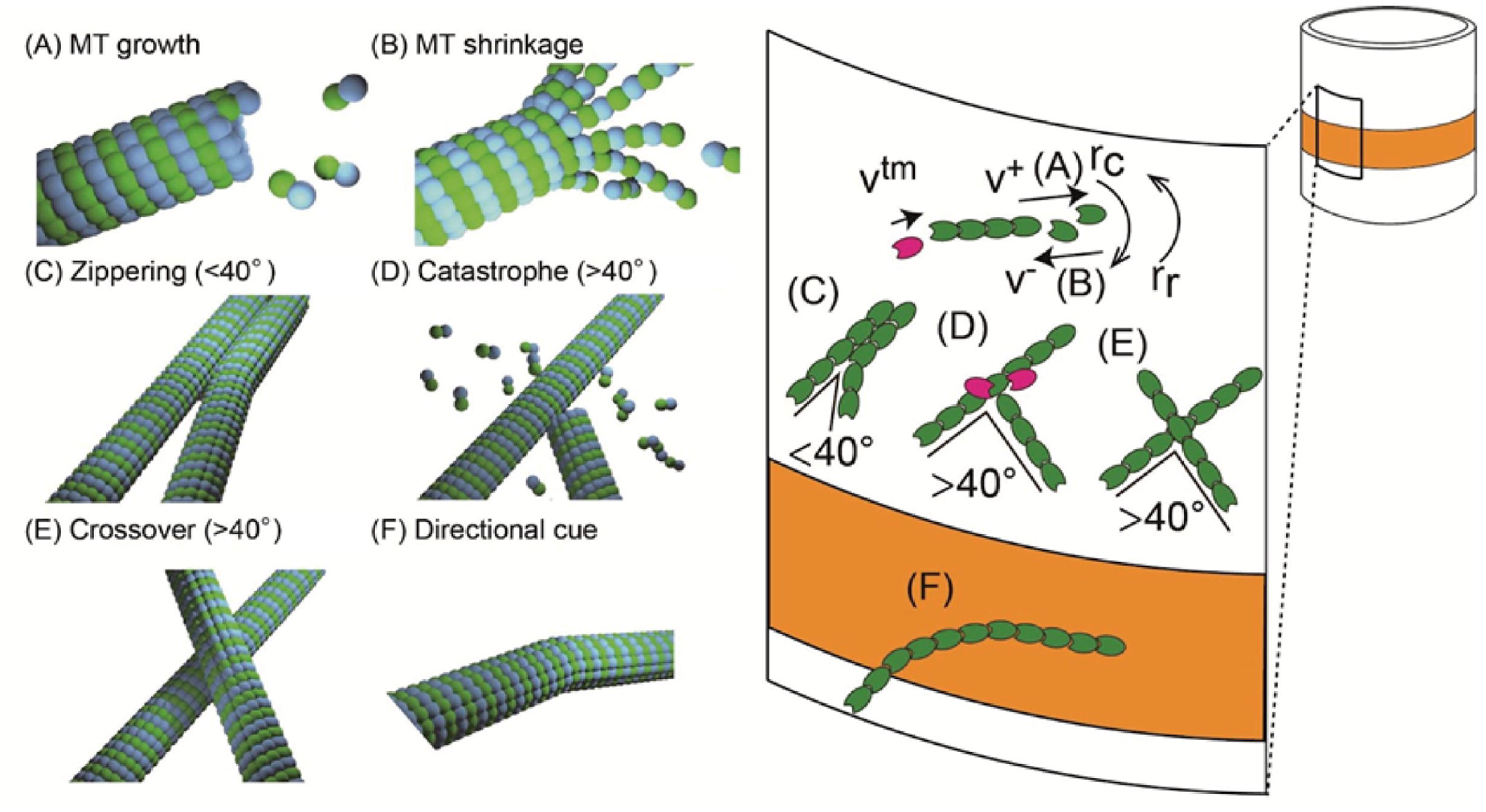
Schematic illustrations of cortical microtubule (CMT) dynamics, their interactions, and the effect of the directional cue (DC). The blue and green spheres in (A–F) represent α and β-tubulins, respectively. The green and magenta oval-like fragments represent the growing and shrinking ends of the microtubule, respectively. The orange region is the DC zone. (A) Microtubule (MT) growth, when the CMT plus end grows at speed *v*^+^. (B) MT shrinkage, when the CMT plus end shrinks at speed *v*^−^ while the CMT minus end undergoes shrinkage at speed *v*^*tm*^. The rescue rate *r*_*r*_, which represents the rate of change from growth to shrinkage at the plus end, and the catastrophe rate *r*_*c*_, which represents the rate of change from shrinkage to growth at the plus end, are also shown. (C) CMT zippering, when one CMT interacts with another within 40° and changes its direction to align with the other. At this angle of interaction, zippering occurs at 100% frequency. (D) Induced CMT catastrophe, when one CMT interacting with another at an angle greater than 40° is induced to depolymerize. (E) CMT crossover, when two CMTs interact with each other at an angle greater than 40° and cross one another. The induced catastrophes (D) and crossovers (E) each occur with a probability of 50% when CMTs meet at an angle greater than 40°. (F) CMTs with a DC, which causes CMTs within the DC zone to change their direction to gradually align to a specific direction (circumferential in this study).

### Interactions of growing CMTs

In addition to the dynamics of a single CMT described above, the interactions between growing CMTs were also implemented in the simulation^22,24,25,26,27^. Three interactions were considered: (1) zippering, when the growing CMT changes the direction of its growing front to be parallel to an existing CMT without changing the direction of the existing CMT, which happens when a growing CMT collides with an existing CMT at an angle smaller than 40° (Fig. 1C); (2) induced catastrophe, when a CMT begins to depolymerize from the growing front when it meets an existing CMT at an angle greater than 40° (Fig. 1D); and (3) crossover, when a growing CMT appears unaffected by the encounter of an existing CMT at an angle greater than 40° (Fig. 1E). For collisions at angles greater than 40°, this model assumes an equal probability (50%) for induced catastrophes and crossovers, based on previous studies^18,25^.

### Implementation of a DC

The length of the CMT agents can be changed, owing to the growth and shrinkage of the CMT plus and minus ends, whereas the orientation of the CMTs does not change by default. With this setup, the DC zone (as shown in orange in Fig. 1F) is the region where the newly added vector (CMT plus end) is biased in a horizontal direction with weight *b*_*d*_:

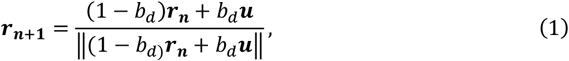

where ***r***_***n***_ and ***r***_***n***+**1**_ are the current and next unit vectors of the CMT plus end, respectively, ***u*** is a circumferential unit vector, and ‖·‖ is the standard norm. The calculation of the unit vector was almost the same as in a previous study^30^, except that random fluctuations in the direction of the CMT growth were not considered in this study. The notations for position, *μ*_*T*_, and width, *σ*_*T*_, of the DC zone along the coordinate z were used (T stands for turn-on). When the input DC zone was moved, the position *μ*_*T*_ was changed computationally at each step. The direction of the newly added CMT plus end depends on the position of the existing plus end, as follows:

(1) When the existing CMT plus end is located outside of the DC zone, a newly added CMT plus end will maintain the original direction.
(2) When the existing CMT plus end is located within the DC zone, a newly added CMT plus end will reorient toward the horizontal direction according to Eq. (1).

### Effect of a finite tubulin pool

To observe CMT dynamics effectively, a finite tubulin pool^27^ was used. The pool limits the CMT length by *L*_*max*_, such that the speed of a growing CMT plus end at any time *t* is dependent on the total length *L*(*t*) of all CMTs in the system:

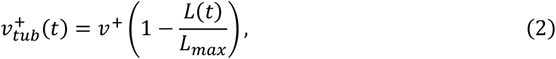

where *L*_*max*_ = *ρ*_*tub*_*A* is the length of all the existing CMTs on the surface area *A* and finite tubulin density *ρ*_*tub*_. In reality, the growth rate of the plus end *v*^+^ can be variable in space; however, it was assumed to be constant in this study.

### Quantification of the width of the CMT band in the data

To quantify the width of the CMT band in the zygote imaging data, cell contours were obtained from the images based on a previous study^34^. Using these cell contours, the cell centerline was extracted and the fluorescence intensity of the CMTs within the cell was projected to the centerline. The fluorescence intensity was fitted with the truncated Gaussian distribution as a function of position z on the centerline using the parameters α, *β, μ, s*, and *b* as follows:

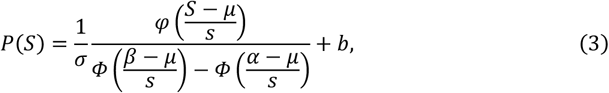

where *S* is the curvilinear coordinate along the cell centerline, and the Gaussian probability density function is 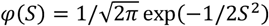 with an average *μ* and standard deviation *s*. The parameter σ is the normalization factor, *b* is the intercept parameter, and *α* and *β* are the lower and upper truncation limits, respectively. The cumulative distribution function is 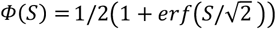. The width of the CMT band is defined as 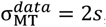, and the CMT position is defined as 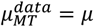. When the band width could not be calculated because the probability distribution became close to a uniform distribution, the zygote was considered to lack a CMT band.

### Quantification of the width of the CMT band in the model

For data analysis, the width of the CMT band can be associated with the probability density of the CMT fluorescence intensity; however, fluorescence intensity was not incorporated into the model, so another way of quantifying the CMT band needed to be defined in the model. For clarity, the term “single CMT formation” was defined as a single CMT oriented in the circumferential direction (a single ring of CMT), and the term “CMT band formation” was defined as a band formed in the circumferential direction (a bundled structure of CMTs). Using these definitions, the width of the CMT band in the simulations was calculated by quantifying the cumulative probability function *P* of the CMT segments. *P*(*S*) was then fitted with a sigmoid curve using five points around *S* = 0.5 (Fig. S1, gray dashed line). The width of the CMT band was estimated as the 68th percentile of the corresponding probability density function for the fitted sigmoidal cumulative distribution (Fig. S1).

### Order parameter tensors quantifying the orientational order of the CMTs

To measure the degree of the orientational order of the CMT array in the simulations, a local order parameter tensor ***q*** and a global order parameter tensor ***Q*** were introduced^27^. Using the orientation of the CMT segment at position *x*, 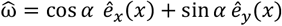, where α ∈ [0, 2π) is the declination angle between 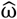 and 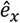. The local order parameter tensor ***q*** at the local area *dA* is defined as

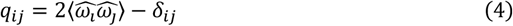

with the matrix form

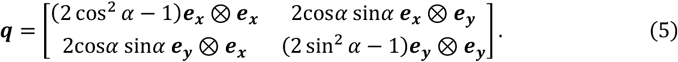

Note that <·> represents the sample average of · within the local area *dA*. The local orientational order and the local orientation of the CMT segments within *dA* were estimated by the first eigenvalue of ***q*** and by the orientation of the corresponding eigenvector of ***q***, respectively. For the characterization of global orientation, the global order parameter tensor was defined as

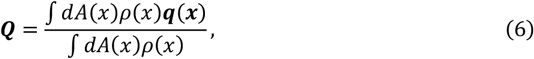

where *ρ*(*x*) is the local areal density of the CMT segments within *dA*. The global orientational order and the global orientation of all the CMTs within the entire region were estimated by the first eigenvalue denoted by *Λ* of ***Q*** and by the orientation of the corresponding eigenvector of ***Q***, respectively. *Λ* becomes 0 for completely random CMT orientations, while for well-organized CMTs, *Λ* becomes +1 for the horizontal array and −1 for the vertical array. The number of local regions was *n*_*div*_ in this study, dividing the whole region into parts with height *H*/*n*_*div*_. A CMT band was considered to exist when Λ exceeded 0.75, and its width was determined from the slope of the probability density distribution.

## Results

### The CMT band moves upward with a speed of 1.5 ± 1.2 μm/min and a width of 5.2 ± 2.4 μm

To determine the mechanism by which the CMT band moves to maintain its position as the zygote expands, we first quantified the width and movement speed of the CMT band using actual observation data (Fig. 2A and 2B). At the CMT-building stage, which occurs just after fertilization, the CMTs are not yet well organized; subsequently, the cell protrudes in the apical direction with the CMT band near the cell tip^1^. The cell maintains anisotropic elongation during the elongation stage, during which the organized CMT band is localized approximately 5–7 µm below the growing tip^2^. To elucidate how the CMT band is maintained, we focused on the elongation stage, assuming a cylindrical cell geometry.

**Fig. 2.**
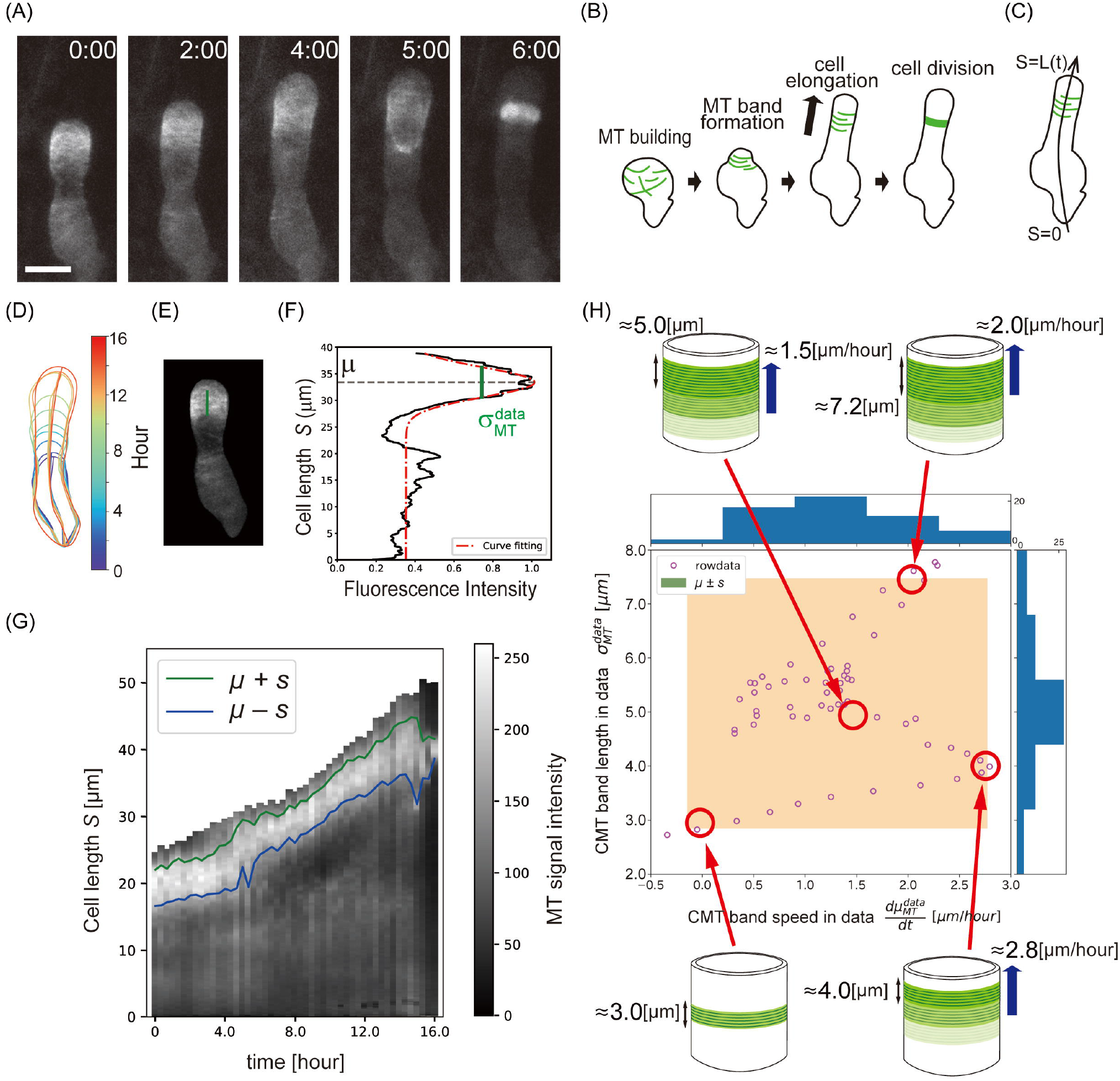
Quantification of the width and movement speed of the cortical microtubule (CMT) band from live-cell imaging data of the Arabidopsis zygote. (A) Two-photon excitation microscopy (2PEM) images of the time-lapse observation of a zygote expressing a MT/nucleus marker based on the previously reported dataset^2^. Numbers indicate the elapsed time (h:min). Scale bar: 10 μm. (B) Schematic illustrations of zygote growth. MT represents microtubule. The black line shows the zygote contour, and green lines show CMT organization. (C) Definition of the cell centerline *S* estimated from the cell contour. (D) Quantification of the cell contour and the cell centerlines. The colors of the contours represent the time from the first frame. (E) Quantification of the CMT band. The green line indicates the width of the CMT band (defined below). (F) Truncated Gaussian fitting of the fluorescence intensity projected on the centerline (black line). The red line represents the fitted truncated Gaussian curve. The position of the CMT band *μ* is estimated based on the median, and the width of the CMT band is estimated as 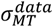. (G) Kymograph of the fluorescence intensity projected on the centerline of the zygote. The green and blue lines indicate the upper and lower limits of the CMT band, respectively, where *s* is the standard deviation. (H) CMT dynamic space (CDS; *n* = 4, indicating the speed and width of the CMT bands with their frequencies [histograms]). The vertical axis denotes 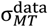, and the horizontal axis is 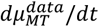. The 95th percentile of the CDS is shown as a pale orange region. Typical CMT movement patterns for the cases shown in the red circles are illustrated schematically from the past band (light green) to the current band (darker green).

To standardize the dynamics of the CMT bands of different individuals to an appropriate time scale, we used the characteristic time *t*_RGS_ at the rapid growth stage (RGS), during which the cell length rapidly increases^2^. We defined the stage from *t*_0_ (5 h before *t*_RGS_) to *t*_RGS_ as the elongation stage. The CMT fluorescence intensity projected on the cell centerline (Fig. 2C and 2D) was fitted by the truncated Gaussian distribution with mean 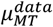 and standard deviation 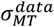 (Fig. 2E and 2F; see Methods). A kymograph of the CMT fluorescence intensity revealed that the CMT band localized to the subapical region, maintaining almost the same distance to the cell tip as the zygote elongated (Fig. 2G). The width of the CMT band also remained constant at ∼5.2 ± 2.4 μm during elongation. The movement speed of the CMT band was less than 2.7 μm/h (Fig. 2H).

### Reorientation strength parameter *b*_*d*_ affects both the width and orientational order of the CMT band

To establish correspondence between the observed data and the model parameters, we introduced a CMT dynamic space (CDS) describing the speed 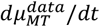 and width 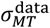 of the CMT band among the previously reported dataset^2^ (Fig. 2H). Eleven parameters were included in the model (Fig. 1): six CMT dynamics parameters, *v*^+^, *v*^−^, *v*^*tm*^, *r*_*r*_, *r*_*c*_, and *r*_*n*_; three mechanical parameters, *b*_*d*_, *μ*_*T*_, and *σ*_*T*_; and two geometrical parameters, *R* and *H* (Table 1). Based on previous studies and the observed data^20,25,27,33^, we set six parameters (*v*^+^, *v*^−^, *v*^*tm*^, *r*_*r*_, *H*, and *r*_*n*_) as fixed and observed the effects of changing the remaining five parameters (*r*_*c*_, *b*_*d*_, *μ*_*T*_, *σ*_*T*_, and *R*). We analyzed the dependence of the five parameters based on the CDS.

To observe the effects of parameter perturbations, it is necessary to identify the parameters that can replicate a similar CMT band with sample-averaged data. We initially fixed the parameter *μ*_*T*_ at the specific location *z* = 10 μm within the range of *z* ∈ [0, 20] (Fig. 2H) and used specific parameters (*r*_*c*_ = 0.001, *b*_*d*_ = 0.01, *σ*_*T*_ = 6.67 [μm], and *R* = 5 [μm]). Using this configuration, we examined the changes in the resulting CMT band by perturbing the parameters (*r*_*c*_, *b*_*d*_, *σ*_*T*_, and *R*).

To avoid misunderstanding, we defined the terms “single CMT formation” as a single CMT oriented in the circumferential direction (a single ring of CMT) and “CMT band formation” as a band formed in the circumferential direction (a bundled structure of CMTs). We noticed that the CMT band could be formed for the parameter *b*_*d*_ = 0, but it was not specifically localized (Fig. 3A). Next, we set *b*_*d*_ = 0.001 for the case of a weak DC, resulting in a CMT band partially localized near the DC zone (dashed line) (Fig. 3B). When *b*_*d*_ = 0.01, for the case of a strong DC, the CMT band covered the entire region of the DC zone (Fig. 3C). We assessed the time scale of the reorientation during single CMT formation. In the case *b*_*d*_ < 0.01, the time required for convergence of CMT bands to the horizontal orientation becomes greater than the time scale of single CMT ring formation, roughly estimated as 2*πR*/*v*_+_ ∼ 6.5 min for *R* = 5 µm (Fig. 3D); therefore, we set the case *b*_*d*_ = 0.01 as a strong DC where the convergence time is fast enough to outpace single CMT ring formation.

**Fig. 3.**
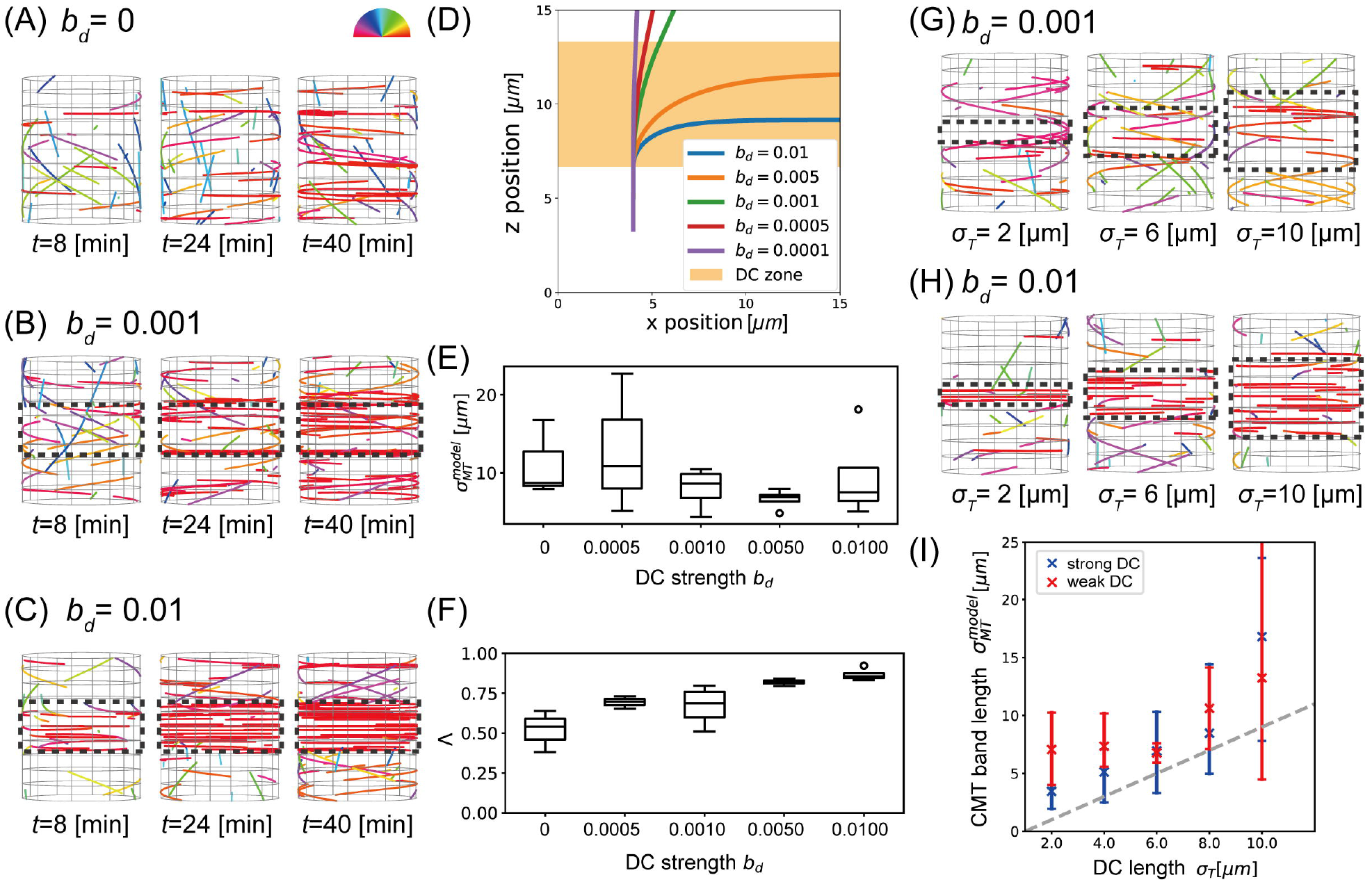
Effect of the tension-responsive ability parameter (*b*_*d*_) and the width of the directional cue (DC) *σ*_*T*_. The boxplots presented below are based on this definition. For all boxplots, the center line (Median): The line inside the box represents the median, which is the middle value of the dataset. Box limits (Upper and Lower Quartiles): The top and bottom of the box correspond to the 75th and 25th percentile, respectively. Whiskers: The whiskers extend to the smallest and largest values within 1.5 times the interquartile range. Points: Data points that fall outside the whiskers are considered outliers and are plotted as individual points. (A–C) Simulation results when using no DC (A), a weak DC (*b*_*d*_ = 0.001; B), and a strong DC (*b*_*d*_ = 0.01; C). The orientation angles of the CMT measured from the horizontal axis are represented by different colors. The DC zone is indicated by black dashed lines. (D) Change in orientation of a single CMT with vertical growth as a function of *b*_*d*_. The DC zone is highlighted in orange. (E) Box plots of CMT width 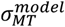 as a function of *b*_*d*_. Box represents the range of the central 50% of data, and small circles represent outliers. (F) The global order parameter *Λ* as a function of *b*_*d*_. (G, H) Results of simulations at *t* = 20 (min) for different *σ*_*T*_ values in the case of a weak DC (G) and a strong DC (H). (I) Width 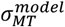 as a function of *σ*_*T*_ in the case of a weak or strong DC. The whiskers extend to the smallest and largest values within 1.5 times the interquartile range.

To quantitatively describe the degree of CMT band formation, we evaluated the width of the CMT band 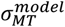 and the global order parameter *Λ*, measuring both the orientation and density of the CMTs (see Methods). The order parameter *Λ* becomes 0 for the completely random orientation of CMTs and 1 for well-organized CMTs. We quantitatively confirmed that the width 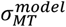 tends to decrease for a certain range of *b*_*d*_, which is consistent with the above qualitative observations (Fig. 3E). Furthermore, we quantitatively confirmed that the order of CMT organization increases as *b*_*d*_ increases, which is also consistent with the observed results (Fig. 3F).

### Catastrophe rate induces a phase transition for both strong and weak DC

Next, to confirm the robustness of the CMT band against CMT depolymerization, we systematically increased the catastrophe rate (*r*_*c*_). Since depolymerization shortens CMTs, it is expected to decrease the probability of CMT collisions that may inhibit the formation of CMT bands. Because the DC may compensate for the increased catastrophe rate by facilitating band formation, we tested the changes for both weak and strong DC conditions. In the case of a weak DC, CMT organization at *t* = 20 min becomes more sparse as the catastrophe rate increases because induced catastrophe occurs frequently (Fig. S2A). Moreover, quantitative evaluation of *Λ* revealed that the degree of orientational order was less than 0.75, indicating that the CMTs are not strongly organized (Fig. S2B). As expected, our definition of the width of the CMT band suddenly dropped to small values at *r*_*c*_ = 0.003, where the phase transition in terms of CMT band width occurs (Fig. S2C). This sudden decrease in the width of the CMT band occurs even in the case of a strong DC, where horizontally aligned single CMT formation is diminished by catastrophe (Fig. S2D). In this case, the index *Λ* was greater than 0.75, indicating that the CMTs are strongly organized (Fig. S2E) and a similar phase transition occurs (Fig. S2F).

### The CMT band widens as the input DC widens under both weak and strong DC

To explore whether the width of the output CMT band 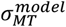 corresponds to the width of the input DC, we focused on the perturbation of the input DC width *σ*_*T*_. In the case of a weak DC, the CMTs at *t* = 20 min were widely distributed beyond the input DC zone (Fig. 3G). In the case of a strong DC, the CMTs at *t* = 20 min were distributed in a region with almost the same width as that of the input DC zone (Fig. 3H). Therefore, we found that the resulting width of the CMT band can be wider than the width of the input DC zone. We quantitatively confirmed that these results were preserved independently of the parameter *σ*_*T*_ (Fig. 3I).

### Cylinder radius does not strongly affect the CMT band but emphasizes two different types of CMT organization

Since changes in the cylinder radius alter the circumference and surface area of the cylinder, which is expected to increase the time required for CMTs to form bands, we tested the cases of *R* = 5.0 μm (typical zygote scale^2^) and *R* = 15 μm. In addition, we simulated weak and strong DC conditions because the DC may compensate for a delay in CMT band formation.

In the case of *R* = 5.0 μm, the orientational order Λ increased in the both the strong and weak DC cases, and the resulting width of the CMT band showed an increasing trend over time (Fig. 4A and 4D). In the case of *R* = 15 μm, the trends were similar, except for the width of the CMT band in the weak DC case. We considered a possible mechanism for this exception as follows.

**Fig. 4.**
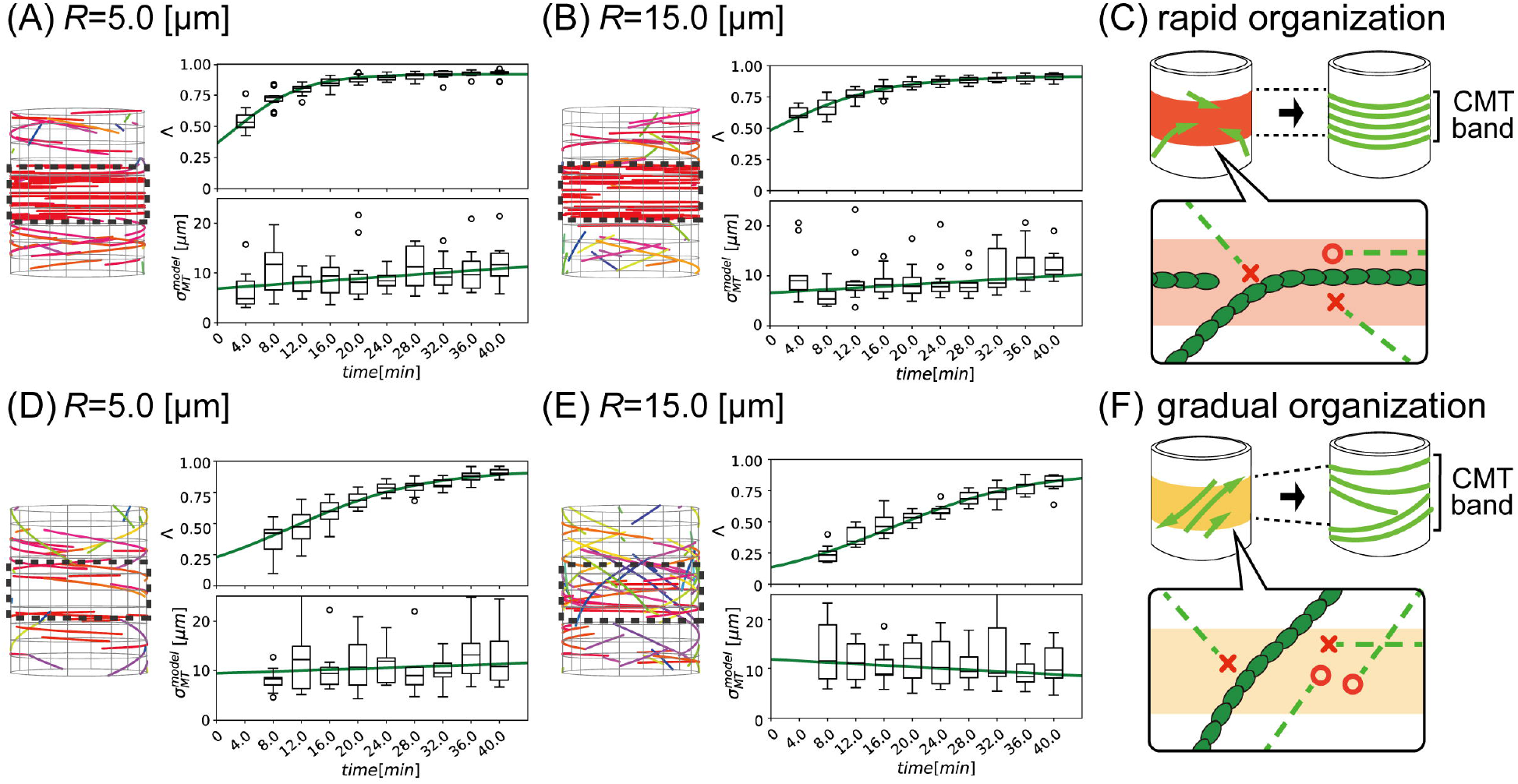
Effect of the cell radius *R* and two different types of cortical microtubule (CMT) organization. (A, B) Simulation results for a strong directional cue (DC) with a temporal dependence of *Λ* and 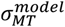 when *R* = 5.0 (μm) (A) and *R* = 15.0 (μm) (B). The green lines represent the sigmoid function for *Λ* and the linear function for 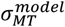. (C) Schematic illustration of rapid CMT organization. Due to the rapid organization of horizontal CMTs, diagonal CMTs may be disorganized, and only the horizontal CMT band remains. The green dashed lines represent newly forming CMTs, where the red circles at the leading edge indicate those that do not collide with pre-existing CMTs, making them likely to grow. By contrast, the red crosses at the leading edge represent CMTs that collide with pre-existing CMTs, indicating a reduced likelihood of growth. The colored band represents the region under strong DC influence. (D, E) Simulation results for a weak DC with a temporal dependence of *Λ* and 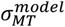 when *R* = 5.0 (μm) (D) and *R* = 15.0 (μm) (E). For cases where the CMT band cannot be defined, the corresponding box plot is not shown. (F) Schematic illustration of gradual CMT organization. The green dashed lines represent newly forming CMTs, where the red circles at the leading edge indicate those that do not collide with pre-existing CMTs, making them likely to grow. By contrast, the red crosses at the leading edge represent CMTs that collide with pre-existing CMTs, indicating a reduced likelihood of growth. The colored band represents the region under weak DC influence. Without horizontal CMTs, diagonal CMTs can be organized so that the CMT band becomes widened.

In the case of a strong DC, the CMT band is rapidly organized, and this behavior is not strongly affected by the radius (Fig. 4A). Therefore, the index *Λ* and the CMT band width *σ*_*MT*_ show a similar trend (Fig. 4A and 4B). By contrast, in the case of a weak DC, the CMTs are not strongly aligned at first and gradually become organized (Fig. 4D and 4E). As a result, the index *Λ* values are lower than those in the case of a strong DC, and the CMT band width *σ*_*MT*_ with *R* = 15 μm gradually decreases, reflecting gradual organization (Fig. 4E).

The mechanism behind this might be the different frequency of CMT encounters, as summarized in Fig. 4C and 4F. In the case of a strong DC, the horizontal CMTs are rapidly organized, preventing the diagonal organization of the CMT band (Fig. 4C). In the case of a weak DC, the horizontal CMTs are not well organized, allowing the diagonal or spiral organization of the CMT band (Fig. 4F). Instead of rapid organization of the CMT band, the newly added CMTs do not quickly reorient but slowly reorient toward the horizontal direction. Consequently, diagonal CMT alignments are formed that do not strongly prevent the vertical organization of CMTs; thus, it takes time for the CMT band to reorganize. We call this phenomenon a gradual organization of CMTs and distinguish this from rapid organization.

### A moving CMT band can form when the DC zone displays an appropriate movement speed and width

As the above results suggest that a desired CMT band can be reproduced with *r*_*c*_ ≤ 0.002, *b*_*d*_ ≥ 0.01, *σ*_*T*_ slightly smaller than the desired band, and *R* = 5.0, corresponding to the observed data, we moved the designated CMT band by changing the specific parameter *dμ*_*T*_/*dt* (the movement speed of the DC zone). As illustrated schematically in Fig. 5A, we searched for the best fitted parameter set (*dμ*_*T*_/*dt, σ*_*T*_) from the CDS in Fig. 2H using an exhaustive search of the model parameters. As the expected DC parameters were unknown, we distributed the parameter set with *dμ*_*T*_/*dt* ∈ [0,4.0] µm/h and *σ*_*T*_ ∈ [0.5,3.5] µm as a similar range to the data.

**Fig. 5.**
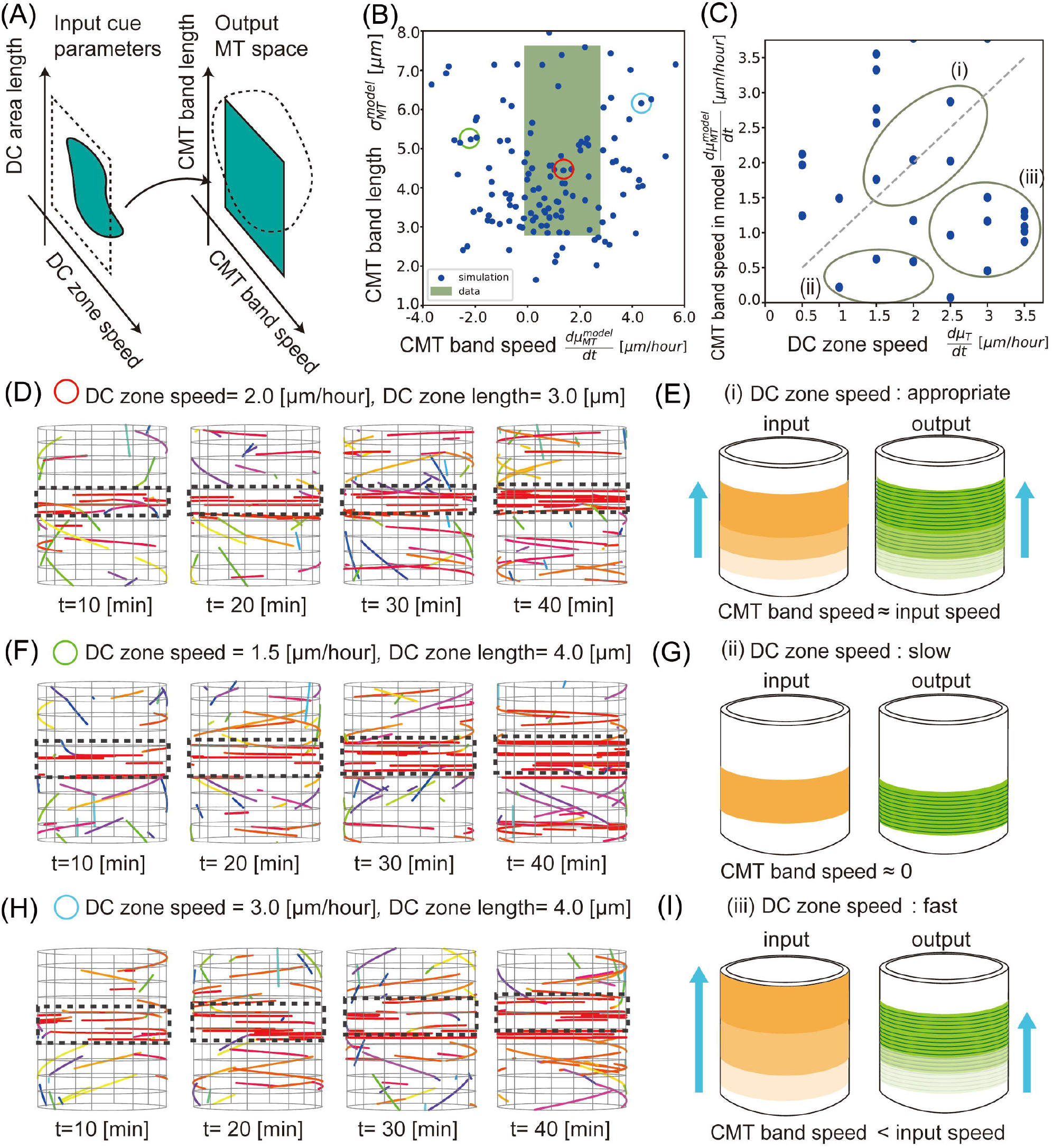
Parameter inference by mapping input directional cue (DC) parameters to the output cortical microtubule (CMT) dynamic space (CDS). (A) Schematic illustrations of the microtubule dynamic (MTD) space in the data, with expected input DC parameters (green) and a similar range of input DC parameters to the MTD space (dashed line). (B) Simulated results in the CDS with the data range from Fig. 2H (green). (C) Discrepancy between the output and input CMT speed compared with the diagonal equivalent line (gray dashed line). Depending on the output speed of the CMT band, there are static states, states that follow the input, and states that lag behind the input. (D, F, H) CMT dynamics with the appropriate DC zone speed and length for the resulting CMT band (Case 01) (D), with a faster speed than the CMT band (Case 02) (F), and with a broader width and faster speed than the CMT band (Case 03) (H). (E, G, I) Schematic illustration of the obtained results. Depending on the input speed of the DC zone, the resulting CMT band did not move (E, i), moved appropriately (G, ii), or lagged behind the DC zone (I, iii).

As a result, 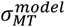 became wider than *σ*_*T*_, while, unexpectedly, the resulting 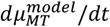 became smaller or sometimes even a negative value, which is below the expected movement speed of the CMT band (Fig. 5B, D, F, H). To demonstrate what happens, we plotted the simulated CMTs from Fig. 5C: Panel (i) shows CMT behaviors with appropriate width and movement speed of the DC zone (Fig. 5D and Fig. 5E), (ii) shows the behavior with slower DC movement (Fig. 5F, G), and (iii) shows the behavior with faster DC movement and a wider DC (Fig. 5H). These results suggest that when the DC zone moves more quickly than the CMT growth speed, the organized CMT band lags behind the DC zone (Fig. 5H and 5I).

In summary, we conclude that a moving CMT band can be achieved with an appropriate width and movement speed of the DC zone.

## Discussion

In this study, we developed an agent-based simulation based on the CMT tension-responsive hypothesis to identify the parameters necessary for the formation of the moving CMT band, which specifically forms in the plant zygote. Through CDS analysis, we successfully compared CMT bands between the model and the observed data to estimate the necessary parameters and obtained four major new findings regarding the reproduction of the moving CMT band in Arabidopsis zygotes: (1) The width of the CMT band is ∼5.2 ± 2.4 μm, and the movement speed is ∼1.3 ± 1.5 μm/h. (2) The output width of the CMT band produced can be wider than the input width of the DC zone. (3) Two types of CMT band formation can occur: rapid organization and gradual organization. (4) The formation of a moving CMT band requires an appropriate width and movement speed of the DC zone.

In addition to the CMT tension-responsive hypothesis, organization of the CMT band has been hypothesized to involve other mechanisms. The edge catastrophe hypothesis, where the cell edge induces CMT catastrophe, was proposed to explain CMT organization in early embryonic apical cells in previous studies^24,25^. However, this hypothesis may not apply to the zygote, especially at the elongation stage, when cell edges are lacking around the CMT band. The conceptual basis for the CMT tension-responsive hypothesis is an increase in the rate of CMT polymerization in the region of higher tension, based on previous studies^12,13^. However, experimental research on CMTs in protoplasts^15^ has shown that tension changes over time may increase the rate of CMT polymerization. Thus, the driving force might be a time derivative of the tension^2^. Additionally, we cannot rule out the possibility that CMTs are severed in a direction perpendicular to the tension exerted by katanin outside the DC zone^8^. Another possibility is an increasing growth rate without reorientation^31^, which is beyond the scope of this study.

To be consistent with the parameter ranges of CMT bands in the actual observed data, the movement speed of the DC zone must be comparable to that of cell elongation (1.5 μm/h on average). The formation of a moving, organized CMT band in the zygote may occur through a similar mechanism to the formation of CMT band structures during the tip growth of fern protonema, whose elongation rate is relatively small^35^. The elongation rate of root hairs is greater than 60 μm/h^36^, which is more than 10 times that of the zygote, which is reported to be 2.4–3.6 μm/h^34^, so it is thought that root hairs are unable to form CMT bands, as supported by our numerical results. One hypothesis is that a “lag behind” phenomenon is associated with tension where the mechanical (tensional) change during elongation becomes so fast that CMT organization does not catch up with the input mechanical perturbation. This might explain why pollen tubes do not have CMT bands, as their elongation speed is more than 10 times faster than that of the zygote^37,38^. Determining when and how organisms evolved a moving CMT band is of great significance, not only for cell and molecular biology, but also for understanding the evolution and diversity of plant cells.

In this study, we developed a data–model correspondence method called CDS to narrow down the range of molecular parameters such as tension-responsive ability and the catastrophe rate of CMTs, which are difficult to observe experimentally. This method can be applied not only to other cylindrical plant cells^39^, but also to cells in general, such as the orientational order during the formation of the extracellular matrix of animal cells (for example, in fruit fly [*Drosophila melanogaster*] trachea^40^). Interdisciplinary research that links biological images and simulation models can thus provide new opportunities for elucidating the essential dynamics of plant and animal cytoskeletons.

## Acknowledgments

The authors thank Koichi Fujimoto, Katsuyoshi Matsushita, and Naoya Kamamoto (Hiroshima University), Takumi Higaki and Haruka Ono (Kumamoto University), Ishimoto Yukitaka (Saga University), and Yusuke Kimata (Tohoku University) for helpful discussions.

## Author Contributions

T.N. and S.T. conceived and designed the study. T.N. and S.T. developed and implemented models and algorithms. H.M., S.N., and M.U. carried out data acquisition from live-cell imaging. T.N. and S.T. wrote the manuscript. T.N., Z.K., S.T., and M.U. edited and reviewed the manuscript. S.T., H.M., and M.U. provided research funding. All authors have read and approved the manuscript.

## Financial Support

This work was supported by the Japan Society for the Promotion of Science [JSPS; KAKENHI (JP20K15832 to S.T.), Grant-in-Aid for Early-Career Scientists (JP22K15135 to H.M.), Grants-in-Aid for Scientific Research on Innovative Areas (JP19H05670 and JP19H05676 to M.U.), Grant-in-Aid for Scientific Research (B) (JP23H02494 to M.U.), International Leading Research (JP22K21352 to M.U.)], the Japan Science and Technology Agency [CREST (JPMJCR2121)], the Suntory Rising Stars Encouragement Program in Life Sciences (SunRiSE; to M.U.), and the Toray Science Foundation (20-6102 to M.U.).

## Data Availability Statement

The data supporting the findings of this study are available from the author, Tomonobu Nonoyama, upon reasonable request.

## Competing Interests

The authors have no competing interests.

## Notes

### Competing Interest Statement

The authors have declared no competing interest.

### Summary of Updates

In this revised version, we have substantially reorganized and rewritten the text to improve clarity and readability. Specifically, we have: Restructured the Introduction to more clearly present the research background and objectives. Revised the Methods section to more accurately describe our experimental procedures Updated several figure legends for better readability. Streamlined the Discussion to avoid repetition and highlight the main findings.

## References

1. Kimata, Y. et al. Cytoskeleton dynamics control the first asymmetric cell division in Arabidopsis zygote. Proc. Natl. Acad. Sci. 113, 14157–14162 (2016).

2. Kang, Z. C. et al. Temporal changes in surface tension guide the accurate asymmetric division of Arabidopsis zygotes, Preprint at 10.1101/2024.08.07.605794 (2025).

3. Heath, I. B. A unified hypothesis for the role of membrane bound enzyme complexes and microtubules in plant cell wall synthesis. J. Theor. Biol. 48, 445–449 (1974).

4. Paradez, A. R., Somerville, C. R. & Ehrhardt, D. W. Visualization of cellulose synthase demonstrates functional association with microtubules. Science. 312, 1491–1495 (2006).

5. Balsuka, F. B., Samaj, J., Wojtaszek, P., Volkmann, D. & Menzel, D. Cytoskelen-plasma Membrane-cell wall continuum in plants. Emerging links revisited. Plant physiol. 133, 482–491 (2003)

6. Lloyd, C. & Chan, J. The parallel lives of microtubules and cellulose microfibrils. Curr. Opin. Plant Biol. 11, 641–646 (2008).

7. Hamant, O. Meyerowitz, E. M., Couder, Y., Traas, J., et al. Developmental patterning by mechanical signals in Arabidopsis. Science. 322, 1650–1655 (2008).

8. Uyttewaal, M. et al. Mechanical stress acts via katanin to amplify differences in growth rate between adjacent cells in Arabidopsis. Cell. 149, 439–451 (2012).

9. Sampathkumar, A. et al. Meyerowitz. Subcellular and supracellular mechanical stress prescribes cytoskeleton behavior in Arabidopsis cotyledon pavement cells. eLife, 3, e01967 (2014).

10. Hervieux, N. et al. Mechanical Shielding of Rapidly Growing Cells Buffers Growth Heterogeneity and Contributes to Organ Shape Reproducibility, Curr. Biol. 27, 3468–3479 (2017).

11. Sapala, A. et al. Why plants make puzzle cells, and how their shape emerges. eLife, 7, e32794 (2018).

12. Inoue, D. et al. Sensing surface mechanical deformation using active probes driven by motor proteins. Nat. Commun. 7, 12557 (2016).

13. Hamant, O., Inoue, D., Bouchez, D., Dumais, J. & Mjolsness, E. Are microtubules tension sensors? Nat. Commun, 10, 2360 (2019).

14. Durand-Smet, P., Spelman, T. A., Meyerowitz, E. M. & Jönsson, H. Cytoskeletal organization in isolated plant cells under geometry control. Proc. Natl. Acad. Sci. 117, 17399–17408 (2020).

15. Colin, L. et al. Cortical tension overrides geometrical cues to orient microtubules in confined protoplasts. Proc. Natl. Acad. Sci. 117, 32731–32738 (2020).

16. Moulia, B., Douady, S. & Hamant, O. Fluctuations shape plants through proprioception. Science. 372, eabc6868 (2021).

17. Dixit, R. & Cyr, R. Encounters between dynamic cortical microtubules promote ordering of the cortical array through angle-dependent modifications of microtubule behavior. Plant Cell. 16, 3274–3284. (2004).

18. Dixit, R. & Cyr, R. The cortical microtubule array: From Dynamics to Organization. Plant Cell. 16, 2546–2552. (2004).

19. Allard, J. F., Wasteneys, G. O. & Cytrynbaum, E. N. Mechanisms of self-organization of cortical microtubules in plants revealed by computational simulations. Mol. Biol. Cell. 21, 278–286 (2010).

20. Eren, E. C., Dixit, R. & Gautam, N. A Three-Dimensional Computer Simulation Model Reveals the Mechanisms for Self-Organization of Plant Cortical Microtubules into Oblique Arrays, Mol. Biol. Cell. 21, 2674–2684 (2010).

21. Deinum, E. E., Tindemans, S. H., Mulder, B. M., Taking directions: the role of microtubule-bound nucleation in the self-organization of the plant cortical array. Phys Biol. 8, 056002 (2011)

22. Shi, X.Q. & Ma, Y-Q., Understanding phase behavior of plant cell cortex microtubule organization. Proc. Natl. Acad. Sci. 107, 11709–11714 (2010).

23. Hawkins, R. J., Tindemans, S. H. & Mulder, B. M. Model for the orientational ordering of the plant microtubule cortical array. Phy. Rev. E, 82, 011911 (2010).

24. Tindemans, S. H., Hawkins, R. J. & Mulder, B. M. Survival of the Aligned: Ordering of the Plant Cortical Microtubule Array. Phys Rev Lett. 104, 058103 (2010).

25. Tindemans S. H., Deinum, E. E., Lindeboom, J. J. & Mulder, B. M. Efficient event-driven simulations shed new light on microtubule organization in the plant cortical array. Front Phys. 2, 1–15 (2014).

26. Chakrabortty, B. et al. A Plausible Microtubule-Based Mechanism for Cell Division Orientation in Plant Embryogenesis. Curr. Biol., 28, 3031–3043. (2018-1).

27. Chakrabortty, B., Blilou, I., Scheres, B. & Mulder, B. M. A computational framework for cortical microtubule dynamics in realistically shaped plant cells. PloS Comput. Biol. 14, e1005959. (2018-2).

28. Schneider, R. et al. Long-term single-cell imaging and simulations of microtubules reveal principles behind wall patterning during proto-xylem development. Nat. Commun. 12, 7085 (2021).

29. Jacobs, B., Saltini, M., Molenaar, J., Filion, L. & Deinum, E. E. Microtubule flexibility, microtubule-based nucleation and ROP pattern co-alignment enhance protoxylem microtubule patterning. Quant. Plant Biol. 6, e2, 1–13 (2024).

30. Mirabet, V., Krupinski, P., Hamant, O., Meyerowitz, E., Jönsson, H., Boudaoud, A. The self-organization of plant microtubules inside the cell volume yields their cortical localization, stable alignment, and sensitivity to external cues. PloS Comput Biol, 14, 21006011 (2018).

31. Li, J., Szymanski, D. B. & Kim, T. Probing stress-regulated ordering of the plant cortical microtubule array via a computational approach. BMC Plant Biol. 23, 308 (2023).

32. Mitchison, T. &’ Kirschner, M. Dynamic instability of microtubule growth. Nature 312, 237–242 (1984).

33. Shaw, S. L., Kamyar, R. & Ehrhardt, D. Sustained microtubule treadmilling in Arabidopsis cortical arrays. Science 300, 1715–1718 (2003).

34. Kang, Z. C. et al. Coordinate normalization of live-cell imaging data reveals growth dynamics of the Arabidopsis zygote. Plant Cell Physiol. pcad020 (2023).

35. Murata, T., Wada, M. Organization of cortical microtubules and microfibril deposition in response to blue-light-induced apical swelling in a tip-growing Adiantum protonema cell. Planta. 178, 334–341 (1989).

36. Grierson, C., Nielsen, E., Ketelaar, T. & Schiefelbein, J. Root hairs. Arabidopsis Book 12, e0172 (2014).

37. Bedinger, P. The remarkable biology of pollen. Plant Cell 4, 879–87 (1992).

38. Schiøtt, M., Romanowsky, S. M., Bækgaard, L. & Harper, J. F. A plant plasma membrane Ca2+ pump is required for normal pollen tube growth and fertilization. Proc. Natl. Acad. Sci. 4101, 9502–9507 (2004).

39. Hasezawa, S., Kumagai, F., Nagata, T., Sites of microtubule reorganization in tabacco BY-2 cells during cell-cycle progression. Protoplasma 198, 202–209 (1997).

40. Öztürk-Çolak, A., Moussian, B., Araújo, S. J., Casanova, J. A feedback mechanism converts individual cell features into a supracellular ECM structure in Drosophila trachea, eLife, 5, e09373 (2016).

