## supplemental figures for "Agent-Based Simulation of Cortical Microtubule Band Movement in Arabidopsis Zygotes": sup_CMT_250404.docx

Supplementary material

Supplementary Figures S1-S2

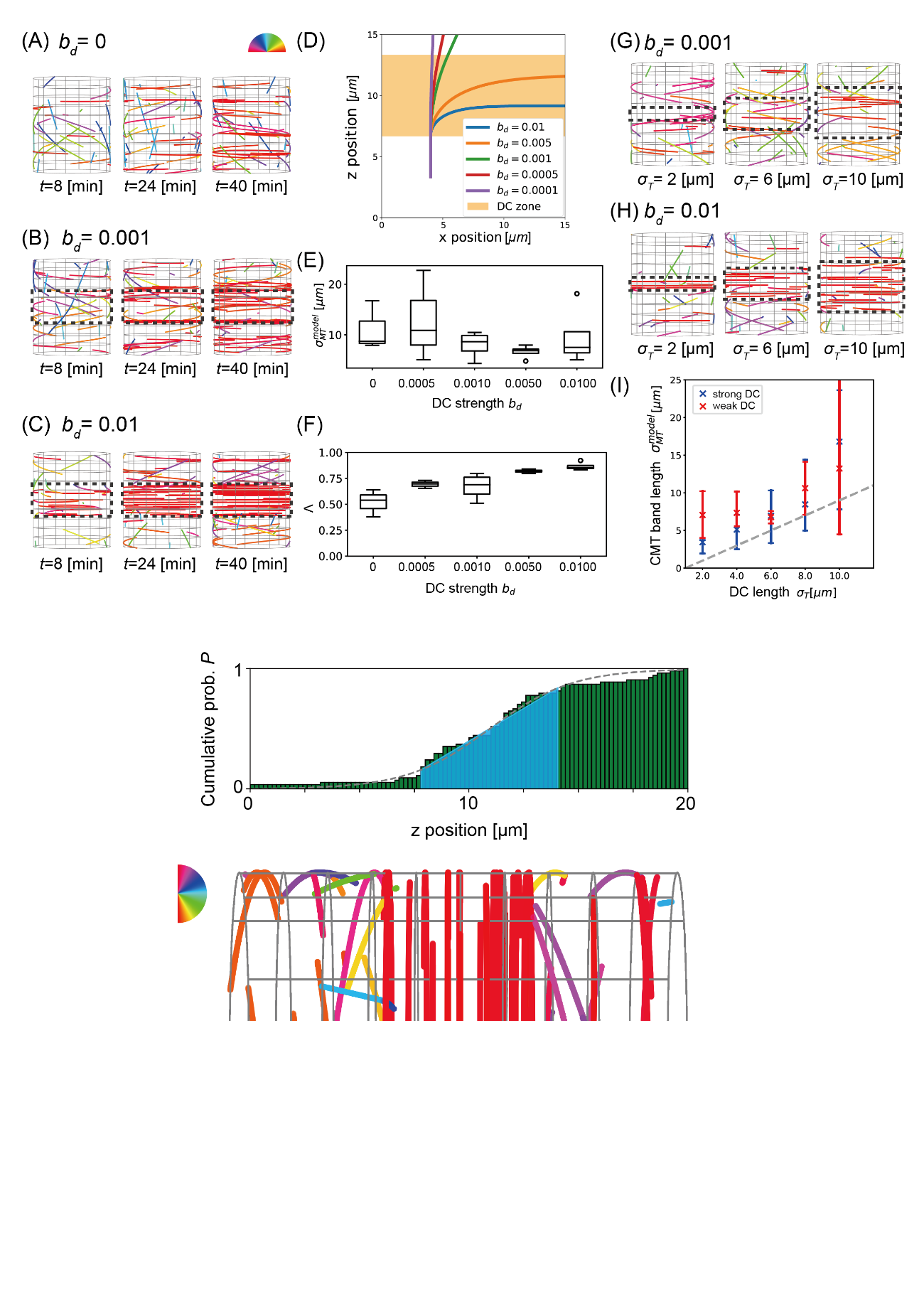

**Supplementary Fig. S1.** The cumulative probability function *P* of the cortical microtubule (CMT) segments in the typical case below. The blue region indicates the 68th percentile of the probability of the CMT, and the gray dashed line indicates the fitted sigmoid curve. The histogram bins correspond to the same area depicted below. The orientation angles of the CMT measured from the vertical axis are represented by different colors.

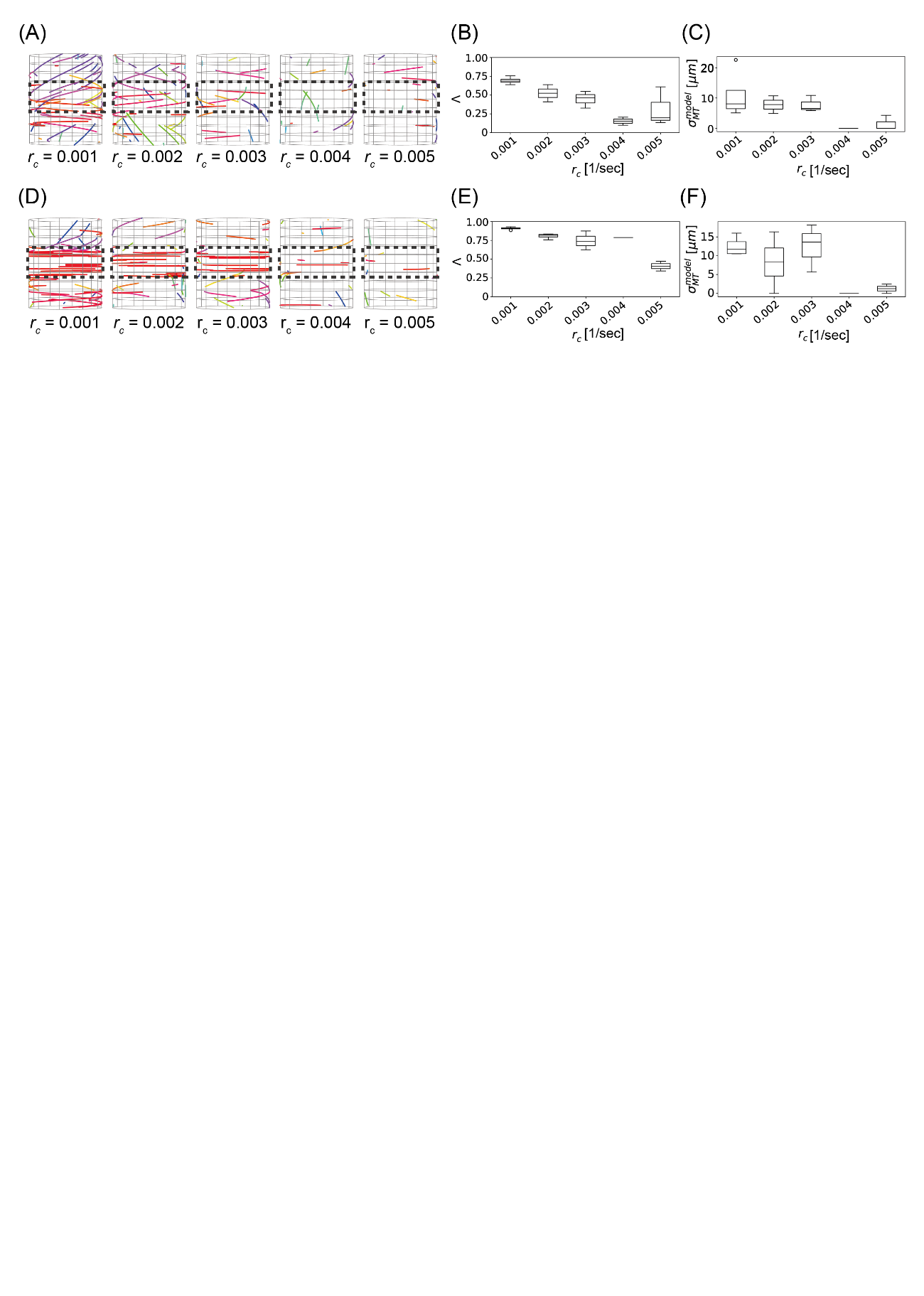

**Supplementary Fig. S2.** Effect of the catastrophe rate $r_{c}$ on the orientational order and width of the resulting cortical microtubule (CMT) band. (A, D) Simulation results at $t = 20$ (min) for different $r_{c}$ in the case of a weak directional cue (DC; A) and a strong DC (D). (B, E) The global order parameter $\Lambda$ as a function of $r_{c}$ in the case of a weak DC (B) and a strong DC (E). (C, F) Width $\sigma_{MT}^{model}$ as a function of $r_{c}$ in the case of a weak DC (C) and a strong DC (F).
